# Larviposition site selection mediated by volatile semiochemicals of larval origin in *Glossina palpalis gambiensis*

**DOI:** 10.1101/802397

**Authors:** Gimonneau Geoffrey, Romaric Ouedraogo, Salou Ernest, Rayaisse Jean-Baptiste, Bruno Buatois, Philippe Solano, Laurent Dormont, Olivier Roux, Jérémy Bouyer

## Abstract

Tsetse flies (*Diptera: Glossinidae*) are the cyclical vectors of trypanosomes, the causative agents of African animal trypanosomosis or nagana in animals and human African trypanosomosis or sleeping sickness in humans. Tsetse flies are K-strategist species with the deposition of a single larva at 10 days intervals in specific sites. As larviposition site selection will strongly impact reproductive success, it is obvious that the selection of larviposition sites is not random and is under strong selective processes, probably mediated by specific cues as suggested by the existence of an aggregation factor in the Morsitans and Fusca groups. This study aimed to highlight the existence of an aggregation effect in the Palpalis group and to test for its chemical nature. We studied the larviposition site selection of *Glossina palpalis gambiensis* according to the presence of conspecific and heterospecific larvae buried in substrates in different settings. Three sets of experiments were performed with either individual or grouped (n = 50) gravid females, and with physical access to substrate or not. In both individual and grouped larviposition experiments, females selected significantly more often trays conditioned by larvae (P<0.005), either conspecific or heterospecific even in the absence of physical contact with the substrate. These results highlight the first evidence for larviposition site selection mediated by volatile semiochemicals of larval origin in *Glossina palpalis gambiensis*. However, these compounds seem not to be species-specific and therefore offer new avenues for the behavioural manipulation of these vectors and for the development of new vector control tools targeting gravid females.

**Author summary:** Larviposition site selection in tsetse flies is govern by several biotic and abiotic factors that lead to an aggregation effect of larvae. Among those, larvae are suspected to produce chemicals that drive females to breeding site but little information is available. This study aimed to highlight the existence of an aggregation effect of larval origin in the Palpalis group and to test for its chemical nature. Through behavioural larviposition choice experiments, we showed that females of *Glossina palpalis gambiensis* deposit their larvae significantly more often in trays conditioned either by conspecific or heterospecific larvae, even in the absence of physical contact with the substrate. These results highlight the first evidence for aggregation effect in *Glossina palpalis gambiensis* mediated by volatile semiochemicals of larval origin. Isolation and identification of these chemicals should offer new avenues for the behavioural manipulation of these vectors and for the development of new vector control tools targeting gravid females.

## Introduction

Oviposition in insects is one of the most important aspects of their reproduction as female breeding site selection can have far-reaching consequences for the life history of the offspring, especially in insects with no parental care (1). To make the best choice for its progeny, females use environmental information mainly driven by olfactory and visual cues. Olfactory cues linked to oviposition are generally volatiles semiochemicals produced by eggs, larvae, habitats, microbes and plants that provide information about the location suitability of breeding sites (2). In hematophagous insects, oviposition pheromones, which mediate interactions between members of the same species, are generally produced by ovipositing female or by conspecific larvae (3). Such compounds are detected by several odour receptors located on the antennae as well as by contact chemoreceptors on tarsi, mouthparts and antennae. Gravid females therefore lay eggs or larvae in suitable breeding sites through information provided by such conspecific cues.

Tsetse flies belong to a unique family in *Diptera* called *Glossinidae*, which is composed of a single genus, *Glossina*. It has been divided into three “groups”, or subgenera, called the Palpalis, Morsitans, and Fusca groups, based on morphological and eco-geographical characteristics. All the 31 species of tsetse flies are theoretically vectors of trypanosomes but only a small number (5 to 6 species) have an epidemiological significance nowadays. They mainly belong to the Palpalis and Morsitans groups of the *Glossina* genus (4). Tsetse flies infest most part of sub-Saharan Africa where they are vectors of African trypanosomoses affecting both humans (HAT: Human African Trypanosomiasis) and livestock (AAT: Animal African Trypanosomosis), two major scourges which impede development in Africa (5). Indeed, HAT is a major neglected tropical disease (6), with more than 50 million people at risk of infection (7) and AAT is considered among the greatest obstacles to the development and intensification of livestock in sub-Saharan Africa (8). There is no vaccine and these diseases lead to the death of patients if left untreated. In the context of WHO’s objectives to eliminate HAT by 2020 (7) and to improve productivity in the African livestock sector to achieve food security (9), innovative one health vector control solutions must be developed.

Tsetse fly control is currently achieved through methods based on insecticide-treated cattle and insecticide-treated attracting devices (traps and targets) and focuses on a unique behavioural target, the “host seeking behaviour” of adults only (10). Other basic life history, behavioural or ecological traits remain largely unexplored despite their importance to tsetse biology and parasite transmission. Although it plays a central role in tsetse population dynamics, larviposition has been one of the most overlooked behaviour.

Compared to other *Diptera*, which basically lay a large number of eggs, the tsetse fly reproductive rate is very low producing a single larva at once. Tsetse flies are larviparous insects and their mode of reproduction is adenotrophic viviparity (11), i.e. the larva develops feeding from the uterine glands of the mother. The first larva is deposited in about 20 days and the subsequent larvipositions occurs at 9 to 10 days intervals thereafter (12). As a result, searching for and choosing a larviposition site is a crucial task for females because the consequences of that choice directly impact their reproductive success and tsetse population dynamics. From an evolutionary point of view, it is obvious that the selection of larviposition sites is not random and is under strong selective pressures to respond to specific cues. Indeed, some physical factors such as shade, colour and soil texture are important for site selection (13,14), but are not specific enough to explain why numerous pupae are found in the same breeding site in the field. The existence of an aggregation factor was suggested in the 1950s (15,16) and an anal exudate produced by larvae have been shown to attract gravid females of the Morsitans group to larval sites (17,18). The chemical compounds present in this exudate is probably not species-specific, which explain field observations of several pupae species at the same breeding site (15).

To date, very few studies have focused on larviposition behaviour, especially in the last twenty years. Nevertheless, there is renewed interest in this behaviour and the use of semiochemicals as attractant for traps or repellent (19,20). Recently, Renda et al. (21) studied the effects of pH, salinity and presence of tsetse pupae in *Glossina brevipalpis* (Fusca group) on larviposition site selection. They showed no effect of pH and salinity, but they observed a larval aggregation effect probably mediated by intra and interspecific chemical cues of larval origin. These results suggest that such an “aggregation factor” could generate similar behavioural responses between flies from different groups and could be used in the field to develop innovative vector control strategies. In West Africa, the main species involve in trypanosomosis transmission belong to the Palpalis groups and there is no information about such “aggregation factor” that may lead to the development of new vector control tools. In this context, this study aims to assess the existence of an aggregation effect in the Palpalis group with a special focus on *Glossina palpalis gambiensis*. We expect that such behaviour is mediated by volatile and/or contact pheromones but this aspect has never been tested yet. Therefore, a special attention will be given to this aspect in this study.

## Materials and methods

### Tsetse flies

All experiments were conducted with *G. p. gambiensis* from a laboratory colony established in 1975 at the Centre International de Recherche-Développement sur l’Elevage en zone Subhumide (CIRDES), Burkina Faso. Flies were maintained under controlled insectary conditions (temperature 25± 2° C, 70± 10% relative humidity and 12:12 h light: dark) and were fed with irradiated bovine blood according to standard rearing procedures using an in vitro silicon membrane feeding system (22). Blood originated from the local abattoir of Bobo-Dioulasso, with whom the CIRDES have a long-term collaboration for blood collection to feed tsetse flies colonies. *Glossina morsitans submorsitans* larvae were also used for substrate conditioning and were obtained from colonized flies in 1979 at the CIRDES and maintained under identical conditions as *G. p. gambiensis*.

In *G. p. gambiensis*, the first larviposition occurs between days 16 and 20 after emergence (15). Therefore, 15 days old gravid females were used for the following experiments.

### Substrate conditioning

White sand commercially available in Bobo-Dioulasso was used as larviposition substrate for all experiments. Sand was washed with tap water and autoclaved at 115° C for 24 hours before being conditioned in autoclaved aluminium trays for larviposition choice experiment. Sand conditioning was achieved by placing 20 freshly deposited larvae of either *G. p. gambiensis* or *G. m. submorsitans* on dry sand. Larvae were allowed to burrow and pupariate into the sand without being removed thereafter.

### Behavioural assays

All experiments were conducted in the insectary under controlled conditions (temperature 25± 2° C, 70± 10% relative humidity and 12:12 h light: dark).

### Experiment 1: Individual bioassays with two choice larviposition sites

In this experiment, individual gravid females *G. p. gambiensis* were offered two choice larviposition sites. Ten experimental cages (30 cm length x 15 cm width x 25 cm height) covered with a net were set up simultaneously. Aluminium trays (25 cm length x 15 cm width x 5 cm height) divided into two equal parts in length (i.e. 12.5 cm) were introduced in each cage (Figure 1). One half received autoclaved sand (control) and the other half received *G. p. gambiensis* conditioned sand (i.e. sand with 20 pupae buried in autoclaved sand). In each cage, one gravid female was released and allowed to larviposit over a five-day period. To reduce mortality and stimulate larviposition, flies were blood fed on the third day using an in vitro silicon membrane feeding system (22). At the end of the experiment, pupae were removed by sifting, counted, and the larviposition site chosen by females was determined. A total of 10 replicates at least were performed per cages and the position of the conditioned sand was randomized between cages.

**Figure 1:**
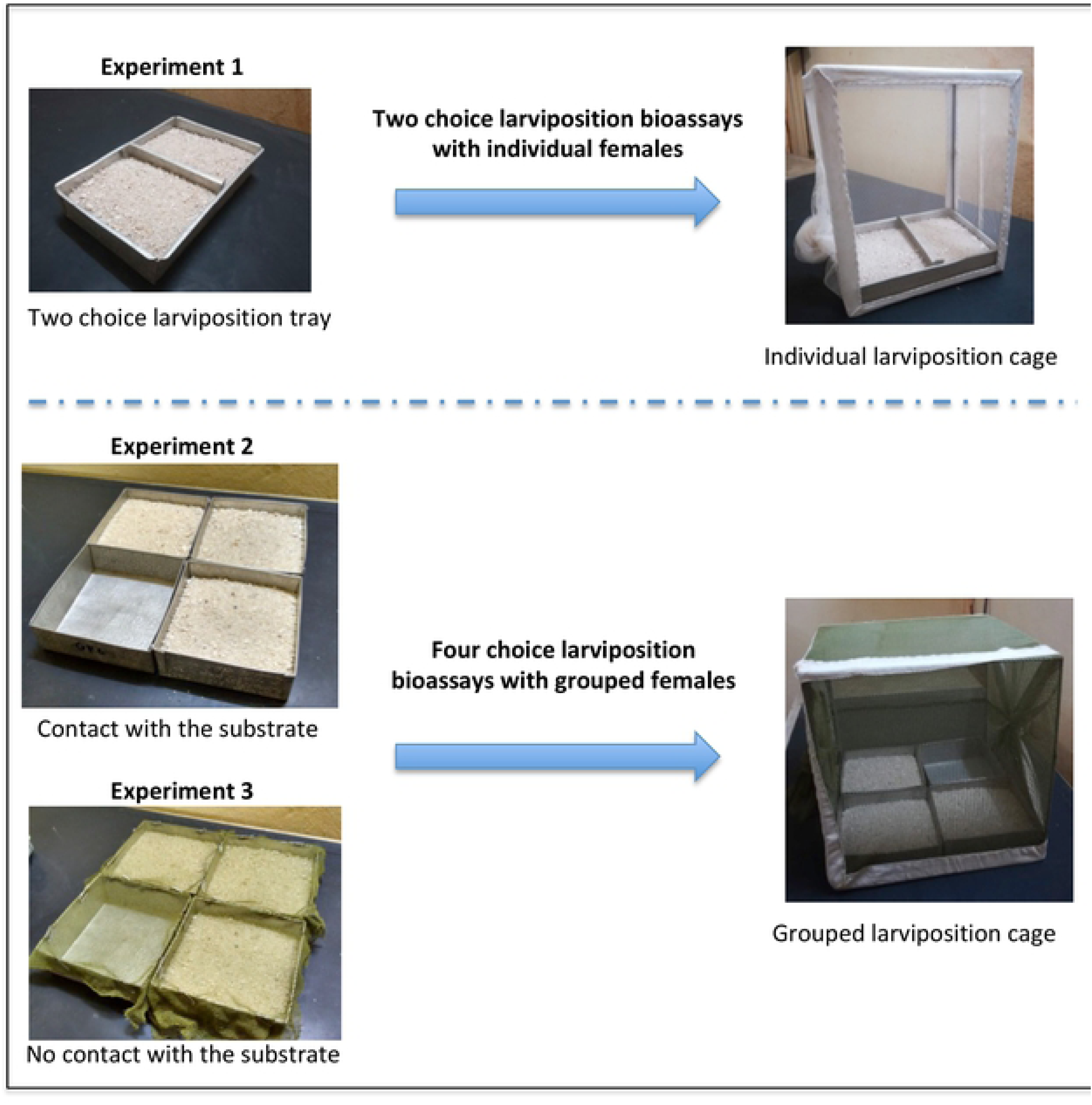
Experimental trays and cages used in experiment 1, 2 and 3.

### Experiment 2: Grouped females bioassays with four choice larviposition sites

In this experiment, four-larviposition sites were proposed simultaneously to 50 gravid *G. p. gambiensis* flies. Four experimental cages (30 cm length x 30 cm width x 30 cm height) were set up simultaneously and ten replicates per cages were performed. The floor of each cage was covered with 4 aluminium trays (15 cm length x 15 cm width and 5 cm height; Figure 1). Two trays received conditioned sand, one conditioned with 20 *G. p. gambiensis* larvae and the other with 20 *G. m. submorsitans.* One of the remaining two trays was kept empty to simulate the larvipositioning conditions in the colony (empty control tray) and the other received only dry autoclaved sand with no pupae (sand control tray). In each cage, 50 gravid females were released and allowed to larviposit over a five-day period after which pupae were removed by sifting and counted. On the third day, flies were blood fed as described previously and in the same time, the sand control and empty control trays were cleaned whereas the two conditioned trays were left undisturbed. The sand control trays were replenished with new autoclaved sand and the possible pupae deposited in the sand and empty trays were collected and numbered. These two unconditioned trays were cleaned on day 3 to prevent additional conditioning by larvae deposited. Larviposition trays were set up following two randomized 4×4 Latin square models among cages. In this set up, flies could be in contact with sand and therefore could use both olfactory and contact cues emanating from conditioned sands to identify the appropriate substrate.

### Experiment 3: Grouped females bioassays with four choice larviposition sites without contact with the substrate

This experiment was identical to the previous one except that physical contacts with sand were not possible, preventing the use of any contact cues to identify the appropriate substrate (the presence of conspecifics). Nets were set up on aluminium trays in order to prevent tsetse flies contact with the substrates. Nets were approximately 2.5 mm diameter allowing larvae to quickly pass through. In these conditions, gravid females could use only volatile cues that emanate from conditioned sands to select larviposition sites. As previously, four experimental cages were set up simultaneously and 10 replicates were performed per cages. Larviposition trays were set up following two randomized 4×4 Latin square models among cages.

### Statistical analyses

In the dual-choice larviposition bioassays, the tray in which the larva was deposited was analysed with a generalized linear mixed model with a binomial family distribution. The location of the conditioned sand in the cage (i.e. in front or behind the cage) and treatments of the sand (conditioned with larvae or autoclaved) and their interactions were used as explanatory variables. The cages and replicates were considered as nested random effects.

In the four-choice larviposition site experiments, the number of larvae deposited was analysed using general linear mixed model with a negative binomial distribution. The larviposition site treatments, site location (i.e. Latin square model) and their interactions were used as explanatory variables. The cages and replicates were considered as nested random effects.

The best models were selected by a stepwise removal of terms, followed by likelihood ratio tests (LRT) (23). Term removals that significantly reduced explanatory power (P <0.05) were retained in the minimal adequate model. The R Software (version 3.1.0) was used to perform all statistical analyses (24).

## Results

### Experiment 1: Individual bioassays with two choice larviposition sites

A total of 106 larvae individually were deposited in either sand conditioned by conspecific larvae or autoclaved sand. Model did not retain the explanatory variable “location of the conditioned sand in the cage” (Likelihood-ratio test: χ^2^ < 0.001, df = 1, *P* = 0.987) but retained the variable “treatment” (Likelihood-ratio test: χ^2^ = 14.97, df = 1, *P*<0.001) showing that *G. p. gambiensis* gravid females deposited significantly more larvae in the conditioned sand with 67 (63.2%) larvae deposited against 39 (36.8%) in autoclaved sand (Table 1).

**Table 1:** Number of larvae deposited in each treatment in the three experiments.

| Treatments | Number of larvae deposited (%) |  |  |
| --- | --- | --- | --- |
|  | Experiment 1 | Experiment 2 | Experiment 3 |
| Control |  | 62 (5) | 237 (18) |
| Sand | 39 (36.8) | 266 (22) | 257 (19) |
| Sand + *G. p. gambiensis* | 67 (63.2) | 493 (40) | 421 (31) |
| Sand + *G. m. submorsitans* |  | 409 (33) | 439 (32) |

### Experiment 2: Grouped females bioassays with four choice larviposition sites

In this experiment, a total of 1230 larvae were deposited by 2000 females, leading to a 61.5% larviposition rate. The total number of larvae deposited at each treatment ranged from 62 (5%) for the control to 493 (40%) for *G. p. gambiensis* treatment (Table 1). The best model retained the larviposition site treatments as significant explanatory variable (Likelihood-ratio test: χ^2^ =226.57, df = 3, *P*<0.001). Gravid females deposit significantly more larvae in sites conditioned with larvae than autoclaved sand alone and control empty trays (Tukey post-hoc test: *P*<0.005 in all cases; Figure 2). However, no differences were observed between sites conditioned either with *G. p. gambiensis* larvae or *G. m. submorsitans* larvae (Tukey post-hoc test value: Z = 1.719, *P* = 0.307; Figure 2). Females were able to differentiate between autoclaved sand and empty trays with significantly more larvae deposited in autoclaved sand (Tukey post-hoc test value: Z = −8.826, *P*<0.001).

**Figure 2:**
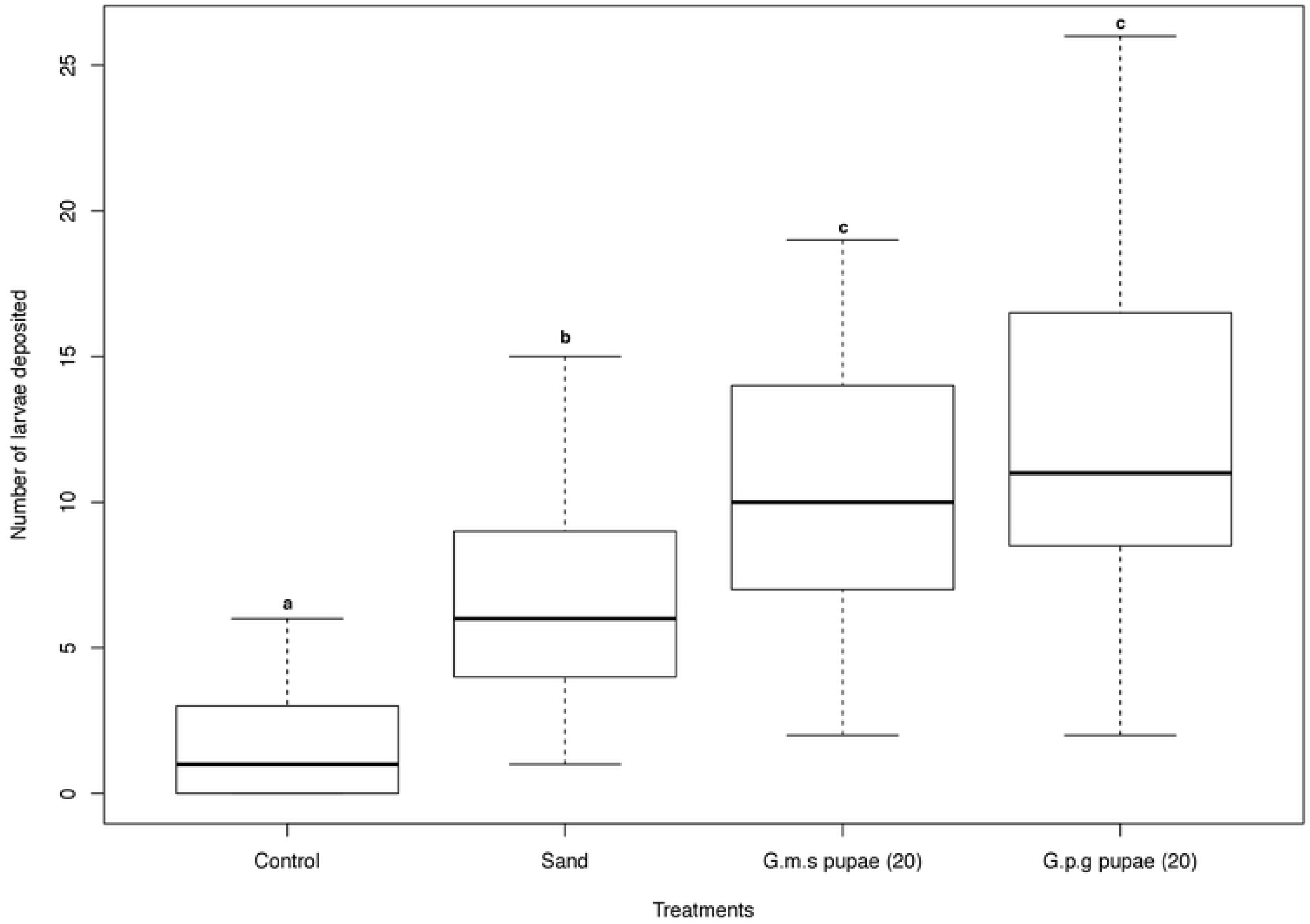
Number of larvae deposited by *G. p. gambiensis* over five days on different substrates: control (empty tray), autoclaved sand and sand conditioned with 20 larvae of either *G. p. gambiensis* or *G. m. submorsitans*. Boxes extend between the 25th and 75th percentile and the thick line denotes the median. The whiskers extend up to the most extreme values. Boxplot with different letters are significantly different (Tukey post-hoc test; P<0.05).

### Experiment 3: Four choice larviposition bioassays for grouped females with no substrate contact

In this last experiment, nets were set upon trays to prevent females contact with the substrate limiting their perception of semiochemicals to volatile compounds only. A total of 1354 larvae was deposited in trays, leading to a 68% larviposition rate. The total number of larvae deposited at each treatment ranged from 237 (18%) for the control to 439 (32%) for *G. m. submorsitans* treatment (Table 1). The best model retained the larviposition site treatments as significant explanatory variable (Likelihood-ratio test: χ^2^ = 26.239, df = 3, *P*<0.001). *G. p. gambiensis* females selected significantly more often sites conditioned with larvae (Tukey post-hoc test: *P*<0.005 in all cases) compared to sand and empty trays, but without being able to make the difference between sites conditioned with *G. p. gambiensis* or *G. m. submorsitans* (Figure 3; Tukey post-hoc test value: Z = −0.282, *P* = 0.992). Without substrate contact, the number of pupae collected from the autoclaved sand site did not differ significantly from that collected from the empty site (Tukey post-hoc test value: Z = −0.508, *P* = 0.957).

**Figure 3:**
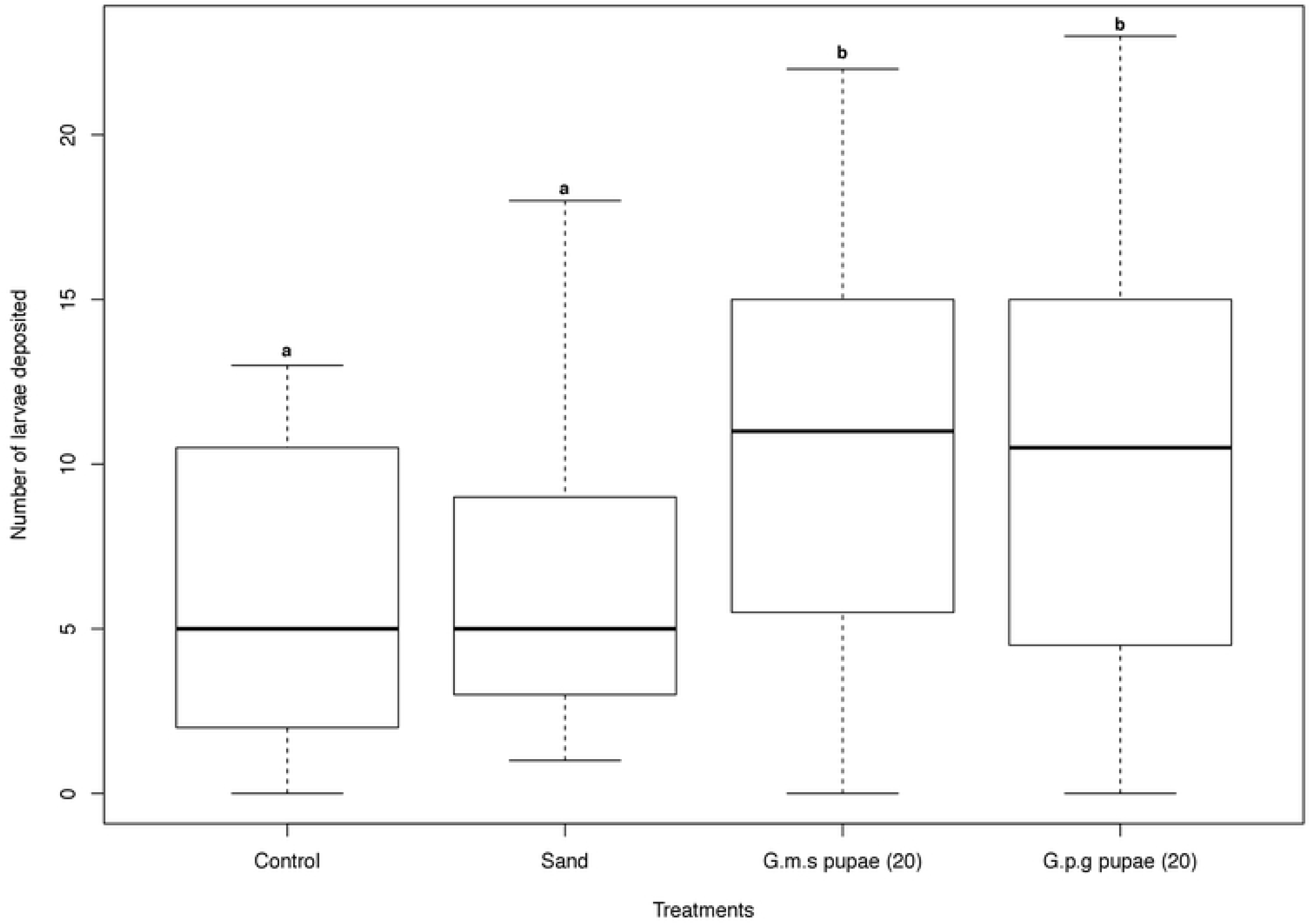
Number of larvae deposited by *G. p. gambiensis* over five days on different substrates: control (empty tray), autoclaved sand and sand conditioned with 20 larvae of either *G. p. gambiensis* or *G. m. submorsitans*. Boxes extend between the 25th and 75th percentile and the thick line denotes the median. The whiskers extend up to the most extreme values. Boxplot with different letters are significantly different (Tukey-post-hoc test; P<0.05).

## Discussion

In this study, we show for the first time a larval aggregation behaviour in tsetse flies of the Palpalis group, *G. p. gambiensis*, mediated by volatile chemicals of larval origin. In both individual and grouped larviposition experiments, females selected significantly more often trays conditioned with either conspecific or heterospecific larvae, even in the absence of physical contact with the substrate. As expected, these results support the hypothesis that larviposition site selection in gravid females is mediated by semiochemicals from larval origin leading to a larval aggregation effect as observed in the field (15,16). Moreover, female discrimination between conditioned and unconditioned tray in absence of physical contact highlights that compounds of larval origin are likely volatile, but did not exclude that contact compounds may occur.

In individual two choice larviposition experiment, females larviposited significantly more often in conditioned substrate (63% larviposition rate). This result indicates that females are able to identify a substrate containing pupae and prefer larvipositing in this substrate rather than in autoclaved sand. Similar results were found by Nash et al. (17) in a grouped two choice larviposition experiment with *G. m. morsitans* with 67% and 63% of larviposition in substrate conditioned with water soluble extracts and ether soluble extracts from larval anal exudates, respectively. Similarly, Leonard and Saini (18) in *G. m. centralis* found that 68% of larvae were deposited in sand conditioned with larvae allowed to pupate and removed before experiment. Therefore, it seems obvious that larvae and/or pupae chemical compounds production acting as an attractant for gravid females is a common evolutionary trait in the three tsetse groups.

In grouped larviposition experiments, gravid females always significantly selected for trays conditioned with larvae, either with substrate contact or not. However, in both experiments, females did not make a choice between conspecific (i.e. *G. p. gambiensis*) or heterospecific (i.e. *G. m. submorsitans*) larvae. In experiment 2, females had access to visual, contact and olfactory cues. In this set up, females did not discriminate between conditioned trays. These results are of primary interest because it suggests that the chemical compounds produced by larvae are not species-specific to the Palpalis and Morsitans groups but rather genus-specific or, at least, present some similarities in their chemical composition. However, these results contrast with a recent study of Renda et al. (21), who showed that *G. brevipalpis* (Fusca group) also selected significantly trays conditioned with larvae but were able to discriminate between conspecific and heterospecific conditioned trays (*G. austeni* larvae, Morsitans group). From a phylogenetic point of view (25), species used by Renda et al. (21) are more distant than those used in the present study. Therefore, it could be suggested that distant species may present more differences in the composition of their aggregation pheromones than closer ones, leading to a better discrimination. However, the selection of the heterospecific conditioned sites as second-best option show that tsetse flies group share interspecific chemical signals related to the aggregation behaviour. As a result, perhaps flies are able to discriminate between both conditioned trays, but they larviposited in both trays because the ecological significance of this message is similar (i.e. appropriate larviposition site in both cases). Moreover, as suggested by Renda et al. (21), a density-dependent factor may also lead females to larviposit in the second best option. If that is the case, it may have hidden a species-specific choice in our study and also could explain field observations of several pupae species found at the same breeding site (15). Therefore, it appears evident that larviposition site selection in tsetse flies, particularly for *G. p. gambiensis*, is mediated by semiochemicals from larval origin. It is however not clear whether each species produces a species-specific or a genus-related pheromone, or even several? Further experiments analysing the chemical composition of such semiochemicals, will help answer these questions.

It is interesting to note that without contact chemoreceptive cues and a limited vision of the substrate due to the net (experiment 3), gravid females were no longer able to differentiate between empty trays and autoclaved sand. From an olfactory point of view, these two trays are probably very similar. Therefore, it could be suggested that vision and contact with the substrate may have important implication in breeding site selection. Indeed, it has been demonstrated in *G. palpalis* that females were able to discriminate soil particle size and prefer rough over smooth surfaces (26) as larvae burrow more readily into rough surfaced soil (14). Moreover, black objects were proven to be attractive as breeding sites (13,26). In our set up, black objects were absent as all larvae burrowed before pupariation but when females had access to substrate (experiment 2), they chose preferentially autoclaved sand as favourable breeding site for their larvae over empty tray. Therefore, contact with the substrate seems to be an important step in the process of larviposition site selection. One could suggest that after 40 years colonization flies would have altered or have lost their larviposition preferences but all these results and others confirm it was not the case (27).

It would be interesting to understand the ecological significance of such a larval aggregation behaviour in tsetse flies. A possible advantage would be for them to subsist at very low density and have extremely low Allee threshold (the density below which a population gets extinct even in the absence of control). It might explain why populations can recover even after strong suppression programs, like for example in Loos islands, Guinea (28), but also in many other places throughout Africa, which may explain many failures in past eradication campaigns (29). The potential implication of this aggregation behaviour on the recently described density dependent dispersal in tsetse should also be investigated (30). Finally, a recently described strategy named “boosted SIT (Sterile Insect Technique)” (31) is currently being tested against tsetse. It is based on the specific transfer of biopesticides, like the hormone analogue pyriproxyfen, from contaminated sterile males to their females. The aggregation behaviour described in the present study may be taken into account as an additional help for this method since it may lead to the contamination of larval sites, as described in mosquitoes (32), which may in turn increase the impact of boosted SIT through the contamination of progeny from other females.

Although significant, our results could have been limited by larval density. Indeed, because there is no information about the carrying capacity of a breeding site before it becomes repellent, it could be suggested that a density dependence factor may act on female behaviour to choose another breeding site. As a result, it can’t be excluded that larvae deposited in heterospecific and unconditioned trays were answers to a density-dependent factor.

In conclusion, this study presents the first evidence for a larval aggregation in *G. p. gambiensis*. Moreover, such behaviour is mediated by volatile pheromones from larval origin but also probably by contact chemoreceptive cues. The ability of tsetse females to recognize and larviposit in conspecific and heterospecific breeding sites suggest tsetse flies’ groups share a similar semiochemicals signature. Isolation and identification of these chemicals should offer new avenues for the behavioural manipulation of these vectors and for the development of new vector control tools targeting gravid females.

## Acknowledgments

We thank the CIRDES insectary technicians for assistance in lab work.

